# Increased HIF-2α Activity in the Nucleus Pulposus Causes Intervertebral Disc Degeneration in the Aging Mouse Spine

**DOI:** 10.1101/2023.12.22.573086

**Authors:** Shira N. Johnston, Maria Tsingas, Rahatul Ain, Ruteja A. Barve, Makarand V. Risbud

## Abstract

Hypoxia-inducible factors (HIFs) are essential to the homeostasis of hypoxic tissues. Although HIF-2α, is expressed in nucleus pulposus (NP) cells, consequences of elevated HIF-2 activity on disc health remains unknown. We expressed HIF-2α with proline to alanine substitutions (P405A;P531A) in the Oxygen-dependent degradation domain (HIF-2αdPA) in the NP tissue using an inducible, nucleus pulposus-specific K19^CreERT^ allele to study HIF-2α function in the adult intervertebral disc. Expression of HIF-2α in NP impacted disc morphology, as evident from small but significantly higher scores of degeneration in NP of 24-month-old K19^CreERT^; HIF-2α^dPA^ (K19-dPA) mice. Noteworthy, comparisons of grades within each genotype between 14 months and 24 months indicated that HIF-2α overexpression contributed to more pronounced changes than aging alone. The annulus fibrosus (AF) compartment in the 14-month-old K19-dPA mice exhibited lower collagen turnover and Fourier transform-infrared (FTIR) spectroscopic imaging analyses showed changes in the biochemical composition of the 14-and 24-month-old K19-dPA mice. Moreover, there were changes in aggrecan, chondroitin sulfate, and COMP abundance without alterations in NP phenotypic marker CA3, suggesting the overexpression of HIF-2α had some impact on matrix composition but not the cell phenotype. Mechanistically, the global transcriptomic analysis showed enrichment of differentially expressed genes in themes closely related to NP cell function such as cilia, SLIT/ROBO pathway, and HIF/Hypoxia signaling at both 14- and 24-months. Together, these findings underscore the role of HIF-2α in the pathogenesis of disc degeneration in the aged spine.

## INTRODUCTION

Intervertebral disc degeneration is one of the most prominent risk factors for chronic low back and neck pain, the leading cause of global disability, and a major contributor to the opioid addiction crisis (1–3). The nucleus pulposus (NP) cells reside at the center of the avascular, and thus hypoxic niche of the intervertebral disc (4,5) and have adapted to this hypoxic environment by controlling the expression and activity of hypoxia-inducible factors (HIFs) (6,7). Two isoforms of HIF-α most prominently expressed in the intervertebral disc are HIF-1α, and HIF-2α that bind to the identical hypoxia response element (HRE) motif and dimerize with a common binding partner HIF-1β/ARNT (8). However, prior studies have demonstrated that there are cell type and tissue type-specific differences between HIF-1α and HIF-2α function and their transcriptomic targets (9,10). A broader categorization of their functions shows that HIF-1α primarily controls cell metabolic activities, whereas HIF-2α regulates cell proliferation, survival, extracellular matrix homeostasis, and oxidative defenses (9). While both HIF-α isoforms are robustly expressed in the NP, HIF-1α is the critical driver of glycolysis, energetics, and overall survival of cells within the specialized niche of the disc (11–14). Notochord-specific *FoxA2^Cre^; HIF-1*α mice show a prominent NP cell apoptosis by birth and subsequently, the NP tissue is replaced by cartilage-like tissue, highlighting the central role of HIF-1α in survival and adaptation of NP cells (15). Noteworthy, in contrast to the dramatic phenotype of *HIF-1α^FoxA2Cre^* mice, our recent investigations pointed to a different function of HIF-2α in the intervertebral disc. *HIF-2*α *^FoxA2Cre^* mice did not result in early cell death, but showed mild and transient protection against age-dependent intervertebral disc degeneration characterized by reduced NP tissue fibrosis (14). While these investigations implied a pro-pathogenic role of HIF-2α in the disc, little is known about the consequences of elevated HIF-2α activity in the NP compartment and its impact on overall intervertebral disc health. This is particularly relevant since increased HIF-2α levels and activity have been seen in human degenerated discs (16). Therefore, the goals of this study were twofold, to establish a causal link between elevated HIF-2α activity and disc health in the aging mouse spine and to identify potential biological pathways responsive to HIF-2α induction in the NP, *in vivo*.

Clinically, humans with *Epas1* gain-of-function syndrome fail to prune or have normal venous regression leading to vascular malformations (17). Notably, mice generated with the homologous *Epas1* mutation showed similar vascular malformations (18). HIF-2α overexpression in adipocytes is shown to promote cardiac hypertrophy (19). Likewise, increased HIF-2 is known to promote cancer cell proliferation (20), and patients with elevated HIF-2 have poor prognosis (21). These data suggest the overall pathological consequences of increased HIF-2 activity in different disease contexts. Concerning musculoskeletal pathologies, in addition to markedly elevated levels in degenerated human discs and enhanced expression of catabolic markers following HIF-2α expression in cultured human NP cells (16), HIF-2α levels are shown to be higher in osteoarthritic human and mouse cartilage, and *Epas1* haploinsufficiency is enough to lessen the severity of traumatic osteoarthritis in mice (22,23). Corroborating these findings; Yang and colleagues showed that induction of HIF-2α levels in knee cartilage of mice with Ad-*Epas1* increased cartilage destruction (24). Some of the effects of HIF-2α on arthritis severity are likely mediated by promoting chondrocyte apoptosis and expression of matrix-degrading enzymes (25). Interestingly, in contrast to these findings, *in vitro* studies of human articular chondrocytes showed the importance of HIF-2α in promoting the expression of healthy cartilage matrix genes (26,27). Likewise, in the Tibetan population, HIF-2α polymorphisms that decrease HIF-2 transcriptional activity and are thus critical in adaptation to a low-oxygen environment, a higher incidence of osteoarthritis and lower back pain has been noted (28,29). Together these results imply that the HIF-2α function in the skeleton is tissue and context-dependent (22–24).

Based on our findings of *HIF-2*α*^FoxA2Cre^* mice, elevated levels seen in degenerated human discs, and its known role in osteoarthritis in mice, we hypothesized that increased HIF-2α expression in the NP would compromise disc health as spine ages. Our results show that elevated HIF-2α activity causes degenerative changes in the NP compartment prominently evident in the aged spine. Our transcriptomic studies showed that elevated HIF-2α activity resulted in the upregulation of genes enriched in biological themes related to primary cilia, cell motility, and axonal guidance by SLIT/ROBO, whereas, genes in themes related to GLP-1, aldolase, and TCA cycle were downregulated reflecting an overall down-modulation of cell metabolism. Supporting our earlier loss-of-function studies in *HIF-2*α*^FoxA2Cre^* mice and based on the findings of this investigation, we conclude that elevated levels of HIF-2α in the NP are one of the pathogenic mechanisms contributing to intervertebral disc degeneration with spine aging.

## METHODS

### Mice

All mouse experiments were performed under protocols approved by the Institutional Animal Care and Use Committee (IACUC) of Thomas Jefferson University following the relevant guidelines and regulations. *Krt19^tm1(cre/ERT)Ggu^*/J (Keratin19^CreERT^ or K19^CreERT^) mice were purchased from Jackson Laboratories. It has been shown K19^CreERT^ allele shows a robust recombination in the NP compartment following tamoxifen injections from P8 up to 15 months of age (30). The *Gt(ROSA)26Sor^tm4(HIF2A*)Kael^*/J (HIF-2α^dPA^, on C57BL6/J background) developed by Dr. William Kaelin were used in the study. HIF-2α^dPA^ (dPA) mice have hemagglutinin-tagged human *HIF2A* cDNA modified with two proline to alanine substitutions (P405A, P531A) introduced 3’ of the mouse *Gt(ROSA)26Sor* promoter in the *HIF-2*α/*Epas1* gene (31) thus promoting the stability of HIF-2α. Control (dPA), homozygous (K19^CreERT^; HIF-2α^dPA^ or K19-dPA) and heterozygous (K19^CreERT^; HIF-2α^dPA/+^ or K19-dPA/+) littermate mice of both sexes were generated and injected with Tamoxifen (3x) at 3 month of age as described before and the phenotype was analyzed at 14- and 24-months (14).

### Histological studies

Spines dissected en-bloc were fixed in 4% PFA for 48 hours at 4°C and decalcified in 20% EDTA at 4°C for 21 days, washed with 1× PBS, and then placed in 70% ethanol before paraffin embedding. Lumbar discs were sectioned in the coronal plane at 7 μm and then stained with 1% Safranin O, 0.05% Fast Green, and 1% hematoxylin. The sections were visualized with Axio Imager 2 microscope (Carl Zeiss) using a 5×/0.15LJN-Achroplan or 20LJ×LJ/0.5LJEC Plan-Neofluar (Carl Zeiss) objective. Images were captured with Axiocam 105 color camera (Carl Zeiss) using Zen2 software (Carl Zeiss). Histological grading of 14 month (*n* = 11 dPA control, 9 K19-dPA/+, 7 K19-dPA) mice, 4-5 lumbar discs/mouse, 33-48 lumbar discs/genotype) and 24-month-old mice (*n* = 9 dPA control, 10 K19-dPA/+, 7 K19-dPA mice, 4-5 lumbar discs/mouse, 33-51 lumbar discs/genotype) was performed by three blinded graders using modified Thompson grading scale and Tessier scale for endplate scoring (32–35). Since the unique interactions between biological, biomechanical, and genetic factors at individual spinal levels have been shown to produce different phenotypic outcomes, each disc was considered as an independent sample (32–34,36).

### Micro-computed tomography (μCT)

μCT imaging was performed on lumbar spines of 14- and 24-month-old mice (*n* = 6 dPA and 6 K19-dPA; 6 lumbar discs and 7 vertebrae/mouse were analyzed) using the high-resolution μCT scanner (SkyScan 1275, Bruker, Konitch, Belgium). Samples were placed in 1× PBS and scanned with an energy of 50LJkV and current of 200 μA, resulting in 15 μm^3^ voxel size resolution. Intervertebral disc height and the length of the vertebral bones were measured and averaged along the dorsal, midline, and ventral regions in the sagittal plane. Disc height index (DHI) was calculated as previously described (14,34,36).

### Picrosirius Red staining and polarized imaging

Picrosirius Red staining and analysis were performed on 7 μm mid-coronal disc sections of 14-month-old mice (*n* = 11 dPA control, 9 K19-dPA/+, 7 K19-dPA, 4-5 lumbar discs/mouse, 33-48 lumbar discs/genotype) and 24-months-old mice (*n* = 9 dPA, 10 K19-dPA/+, 7 K19-dPA , 4-5 lumbar discs/mouse, 33-51 lumbar discs/genotype) as described before (37). The stained sections were imaged using Eclipse LV100 POL (Nikon, Tokyo, Japan) with a 10×/0.25 Pol/WD 7.0 objective and Nikon’s Digital Sight DS-Fi2 camera. Under polarized light, stained collagen bundles appear as either green, yellow, or red pixels that correlate to fiber thickness indicating thin, intermediate, or thick fiber thickness, respectively (38). Using NIS Elements Viewer software (Nikon, Tokyo, Japan), fibers were quantified by thresholding for green, yellow, and red pixels over the selected region of interest (ROI) for NP and AF.

### Immunohistopathology and digital image analysis

Mid-coronal disc sections (7 μm) were deparaffinized in histoclear and rehydrated in graded ethanol solutions (100-70%), water, and 1× PBS. Antigen retrieval was performed using either citrate-buffer, proteinase-K, or MOM kit (Vector Laboratories, Burlingame, CA, USA; BMK-2202) depending on the antibody. Sections were blocked for 1 hour, at room temperature, in 5% to 10% normal goat or donkey serum as appropriate in PBS-T (0.4% Triton X-100 in PBS) or using the reagent from the MOM kit. Then, they were incubated at 4°C overnight in primary antibodies against anti-HA (1:100; BioLegend; 901533), CA3 (1:150; Santa Cruz Biotechnology; sc-50715), CS (1:300; Abcam; ab11570), ACAN (1:50; Millipore; ab1031), ARGxx (1:200; Abcam; ab3773), COMP (1:200, Abcam; ab231977), COL I (1:100; Millipore; abt123), COL II (1:400; Fitzgerald; 70R-CR008), After incubation with primary antibody, tissues were washed and reacted with Alexa Fluro-594 (Ex: 591 nm, Em: 614 nm) conjugated secondary antibody (Jackson ImmunoResearch Laboratories, West Grove, PA, USA) at a 1:700 dilution in blocking buffer for 1 hour at room temperature. The sections were then washed and mounted with ProLong Gold Antifade Mountant with DAPI (Thermo Fisher Scientific, P36934). Slides were visualized with the Axio Imager 2 (Carl Zeiss Microscopy) using 5×/0.15LJN-Achroplan (Carl Zeiss Microscopy) or 10×/0.3 EC Plan-Neofluar (Carl Zeiss Microscopy) or 20×/0.5 EC Plan-Neofluar objectives, X-Cite 120Q Excitation Light Source (Excelitas Technologies), AxioCam MRm camera (Carl Zeiss Microscopy), and Zen2 software (Carl Zeiss Microscopy). All quantifications were conducted in 8-bit grayscale using the Fiji package of ImageJ (39). Images were thresholded to create binary images, and NP and AF compartments were manually segmented using the Freehand Tool. These defined regions of interest were analyzed either using the Analyze Particles (cell number quantification) function or the Area Fraction measurement.

### FTIR imaging spectroscopy and spectral clustering analysis

FTIR spectral acquisition and analysis utilizing deparaffinized sections (7 μm) of decalcified mouse lumbar disc tissues from control and mutant animals at the 14- and 24-months (*n* = 6 mice/genotype/age, 2 disc/mouse, 12 total discs/genotype) were performed as described previously (34,36). Spectrum Spotlight 400 FT-IR Imaging system was used to acquire spatial resolution images in the mid-IR region from 4000 to 800 cm^−1^ (Perkin Elmer, Waltham, MA, USA) of three consecutive sections/disc to minimize section-based variation. ISys Chemical Imaging Analysis software (v. 5.0.0.14) was used to analyze the mean second-derivative absorbances which were quantified and compared between the genotypes. Additionally, collected spectra were analyzed using *K*-means clustering analysis in the Eigenvector Solo+MIA software (v. 8.8) to agnostically delineate anatomical regions within the disc. Regions of IR images are separated into two or more classes, or “clusters,” according to spectral similarity. The *K*-means in our analyses was *K* = 7. During each iteration, the remaining objects (pixels of the spectral image) are assigned to one of these clusters based on the distance from each of the *K* targets. New cluster targets are then calculated as the means of the objects in each cluster, and the procedure is repeated until no objects are reassigned after the updated mean calculations.

### Tissue RNA isolation and Microarray analysis

NP tissues were microdissected from 14M and 24M control dPA and K19-dPA mice. Tissues from Ca1/2-Ca12/13 from each mouse were pooled and stored into RNAlater Reagent (Invitrogen, Carlsbad, CA, USA) at -80°C till isolation (*n* = 6 mice/genotype). Samples were homogenized with a Pellet Pestle Motor (Sigma Aldrich, St. Louis, MO, USA; Z359971), and RNA was extracted using RNeasy Mini Kit (Qiagen, Valencia, CA, USA). DNA-free RNA was quantified on a Nanodrop ND-100 spectrophotometer (Thermo Fisher Scientific, Waltham, MA, USA) and quality was assessed using Agilent 2200 TapeStation (Agilent Technologies, Palo Alto, CA, USA). Using GeneChip WT Plus kit, the fragmented biotin-labeled cDNA was synthesized from the extracted RNA according to ABI protocol (Thermo Fisher Scientific) and hybridized to GeneChips (Mouse Clariom S). After washing and staining with GeneChip hybridization wash and stain kit, the chips were scanned on the Affymetrix Gene Chip Scanner 3000 7G using Command Console Software. Expression Console Software v 1.4.1 was used to perform quality control and to generate CHP files by sst-rma normalization from Affymetrix CEL file. Affymetrix Transcriptome array console 4.0 software was used to perform hierarchical clustering of samples and to determine DEGs, only protein-coding genes with p ≤0.05 and fold change of ≥1.7 were considered. We then used “COMPBIO (COmprehensive Multi-omics Platform for Biological InterpretatiOn) to understand the functional relevance of the DEGs. The platform can analyze single or multi-omic data entities (genes, proteins, microbes, metabolites, miRNA) and deliver a holistic, contextual map of the core biological concepts and themes associated with those entities. The array data generated and analyzed during this study is available in NCBI Gene Expression Omnibus (GEO) repository (GSE249908).

### Statistical analysis

Statistical analysis was performed using Prism 9 (GraphPad, La Jolla, CA, USA) with data presented as whisker box plots showing all data points with median and interquartile range and maximum and minimum values. Differences between distributions were checked for normality using Shapiro-Wilk tests and further analyzed using an unpaired *t* test (two-tailed) for normally distributed data and the Mann-Whitney *U* test for non-normally distributed data. Comparison of multiple groups of normally distributed data was performed using one-way ANOVA, whereas non-normally distributed data were analyzed using Kruskal-Wallis and Dunn’s or Welch’s multiple comparison tests. Analyses of Modified Thompson Grading distributions and fiber thickness distributions were performed using a chi-square test at a 0.05 level of significance. Since the unique interactions between genetic, biological, and biomechanical factors at individual spinal levels have been shown to produce different phenotypic outcomes in humans and mice, each disc was considered an independent sample (32,36,40–42).

## RESULTS

### Elevated HIF-2**α** activity impacts NP compartment morphology in aged mice without affecting cell survival

We have previously shown that the deletion of HIF-2α in the NP compartment (*HIF-2α^FoxA2Cre^*) causes transient protection from early degenerative changes in 14-month-old, middle-aged mice (14). We therefore hypothesized that elevated HIF-2α levels and activity in NP will accelerate age-dependent degenerative changes. To test this hypothesis, we conditionally expressed HIF-2α (HIF-2αdPA) in NP that is resistant to prolyl hydroxylase-mediate degradation and studied the disc phenotype of mice as a function of spine aging. Accordingly, *HIF-2_α_^dPA^* mice were crossed with an NP-targeting tamoxifen-inducible K19^CreERT^ driver, and the resulting littermate dPA control, K19-dPA/+, and K19-dPA mice were injected at 3-months (3M) and aged up to 24-months (24M) (Fig. 1A). To confirm the expression of HA-tagged HIF-2αdPA protein in the disc, we stained the disc sections with an anti-HA antibody which showed cellular staining in the NP compartment of K19-dPA but not in the control dPA mice (Fig. 1B).

**Figure 1.**
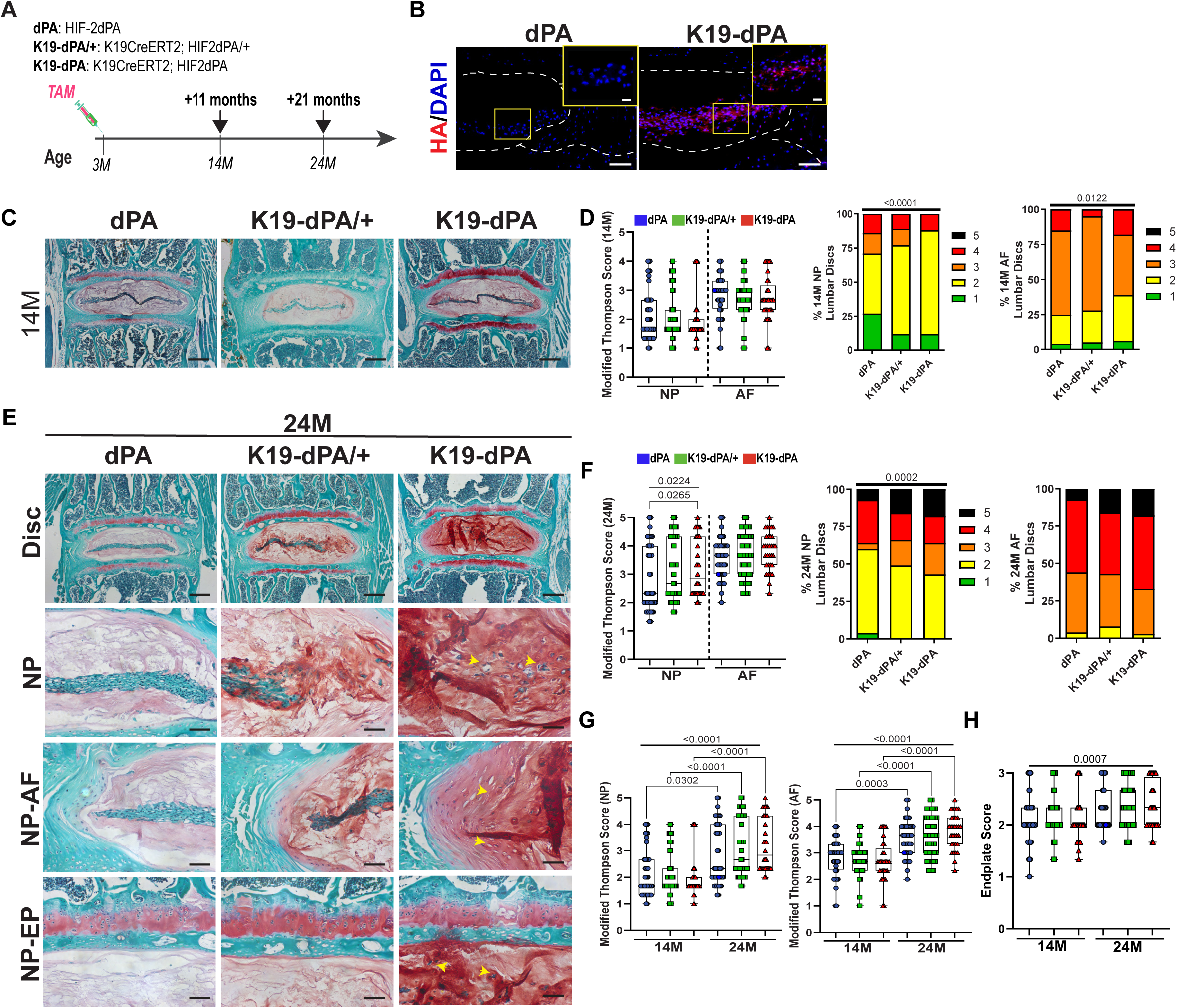
Elevated HIF-2α levels in the NP cause age-dependent disc degeneration in mice. (A) Timeline of experimental mice generation and analysis. (B) K19-dPA mice show expression of HA-tagged HIF-2dPA in the lumbar discs of 14-month (14M) dPA and K19-dPA mice stained with anti-HA antibody, n = 2 discs/animal, 3 animals/genotype (scale bar = 100 μm; insert scale bar = 25 μm). (C) Safranin O/Fast Green staining of lumbar discs showing disc morphology and overall proteoglycan content in the intervertebral disc in 14M control (dPA), heterozygous (K19-dPA/+), and homozygous (K19-dPA) animals (scale bar = 200LJμm). (D) Histological grading assessment of 14M lumbar using the modified Thompson scale. (E) Safranin O staining of lumbar discs in 24M animals. Yellow arrowheads indicate the loss of cell band and fibrotic remodeling in the NP along with the loss of demarcation between NP and AF compartments. Scale bars, top row: = 200μm, bottom rows: 50μm (F) Histological grading assessment of NP and AF using the modified Thompson scale. (G) Comparison of histological grading scores as a function of aging within each genotype. (H) Endplate grading assessment at 14M and 24M. *n* = 14M: 11 dPA, 9 K19-dPA/+, 7 K19-dPA mice, 4-5 lumbar discs/mouse, 33-48 lumbar discs/genotype and 24M: *n* = 9 dPA, 10 K19-dPA/+, 7 K19-dPA mice, 4-5 lumbar discs/mouse, 33-51 lumbar discs/genotype. The significance for grading distribution was determined using a chi-square test. The significance of differences among 3 or more groups was determined using one-way ANOVA or Kruskal–Wallis with Dunn’s test. Quantitative measurements represent the median with the interquartile range.

To understand if increased HIF-2α levels impact intervertebral disc morphology, sections were stained with Safranin O, Fast Green, and Hematoxylin. Histological assessment of 14-month-old (14M) mice showed no apparent differences in NP and AF compartment morphology, and overall disc structural integrity was comparable between the genotypes with no signs of disc herniation (Fig. 1C). At this age, a majority of discs, regardless of the mouse genotype showed vacuolated NP cells surrounded by proteoglycan-rich extracellular matrix. The endplate appeared normal, and the AF showed concentric lamellae interspersed with fibrocartilaginous AF cells. However, as expected, NP and AF compartments of some discs showed signs of early degeneration with smaller NP cell band size and distortion of AF lamellae along with disruption of the NP-AF tissue interface. However, these aging-associated changes were present across all the genotypes. Accordingly, at 14 months, the NP and AF average grades between genotypes were comparable, and the grade distributions showed only small differences with mutant mice showing a slight reduction in the percentage of healthy grade 1 discs (Fig. 1D). At 24 months (24M), however, there was a small but significant increase in NP average grades of degeneration in K19-dPA mice (Figure 1E). This was also captured by the changes in the grade distributions showing a pronounced increase in the percentage of highly degenerated grade 5 discs with severe annular defects and contained herniations (Figure 1E). When the grading data within each genotype was compared across the ages, the increase in average grades of degeneration was more pronounced in K19-dPA mice compared to dPA control animals in both the NP and AF compartments (Fig. 1F-G). Scoring of the cartilaginous endplates (EP) using the Tessier grading scale (32) showed no discernable differences in EP scores across the age and genotype (Fig. 1H).

To assess if there were corresponding changes in disc hydration-related parameters, microCT measurements of disc height (DH), and vertebral body height (VBH), were performed. At both 14M and 24M, K19-dPA mice showed no differences in DH or VBH. However, when disc height measurements were normalized with vertebral lengths to calculate the disc height index (DHI), at 14M, there was a small but significant decrease in DHI compared to control mice which was attenuated as the mice aged to 24M (Figure 2A-B). These analyses showed that elevated expression of HIF-2α in NP cells may promote mild degenerative changes in the disc and affect disc hydration.

**Figure 2.**
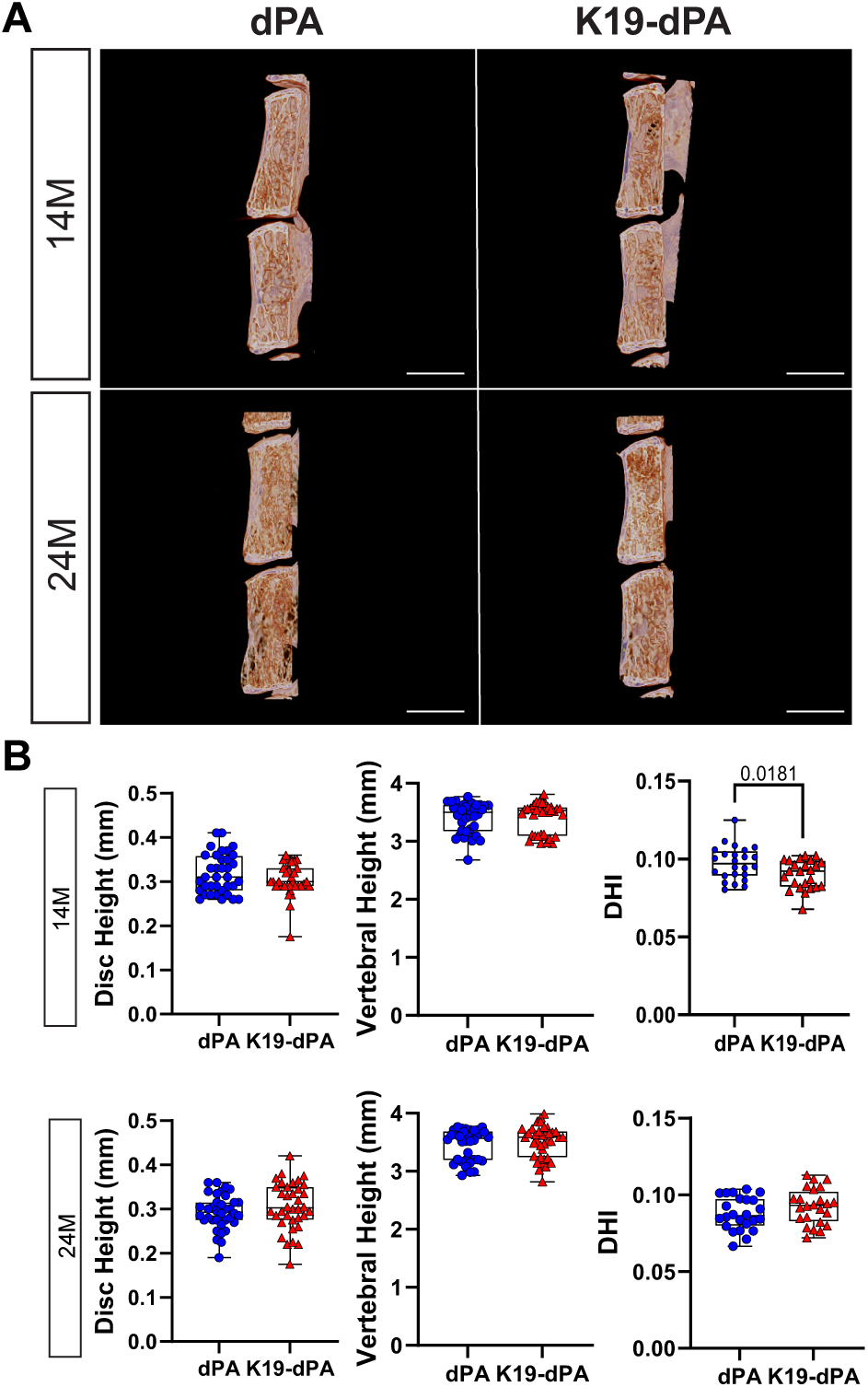
Increased HIF-2α activity in NP results in a decreased disc height index. **(A)** Representative reconstructed μCT images of hemi-sections of lumbar motion segments in 14- and 24-month-old dPA and K19-dPA mice, scale bar = 1mm. Lumbar disc height, vertebral height, and disc height index (DHI) in (B) 14M and (C) 24M dPA and K19-dPAmice. *n* = 14M: 6 dPA, 6 K19-dPA mice; 24M: 6 dPA, 6 K19-dPA mice; 6 lumbar discs and 7 vertebrae/mouse were analyzed. Significance for quantitative measures was determined by using Mann–Whitney *U* test or an unpaired t test with Welch’s correction, as appropriate. Quantitative measurements represent the median with the interquartile range.

### K19-dPA mice show changes in AF collagen turnover and disc matrix chemical composition

To understand if the overexpression of HIF-2dPA in the NP affected collagen turnover in the AF compartment, the organization of the collagen matrix was assessed using Picrosirius Red staining coupled with polarized light microscopy. Polarized imaging revealed that at 14M, there were fewer thin and more intermediate fibers in the AF indicating less collagen turnover in the AF compartment of K19-dPA mice (Figure 3A-C). However, at 24M, the differences noted between the genotypes in both quantity and distribution at 14M were not apparent. These analyses suggest that increased HIF-2α in the NP affects AF collagen turnover during early aging but that these changes are secondary to the effects of spine aging.

**Figure 3.**
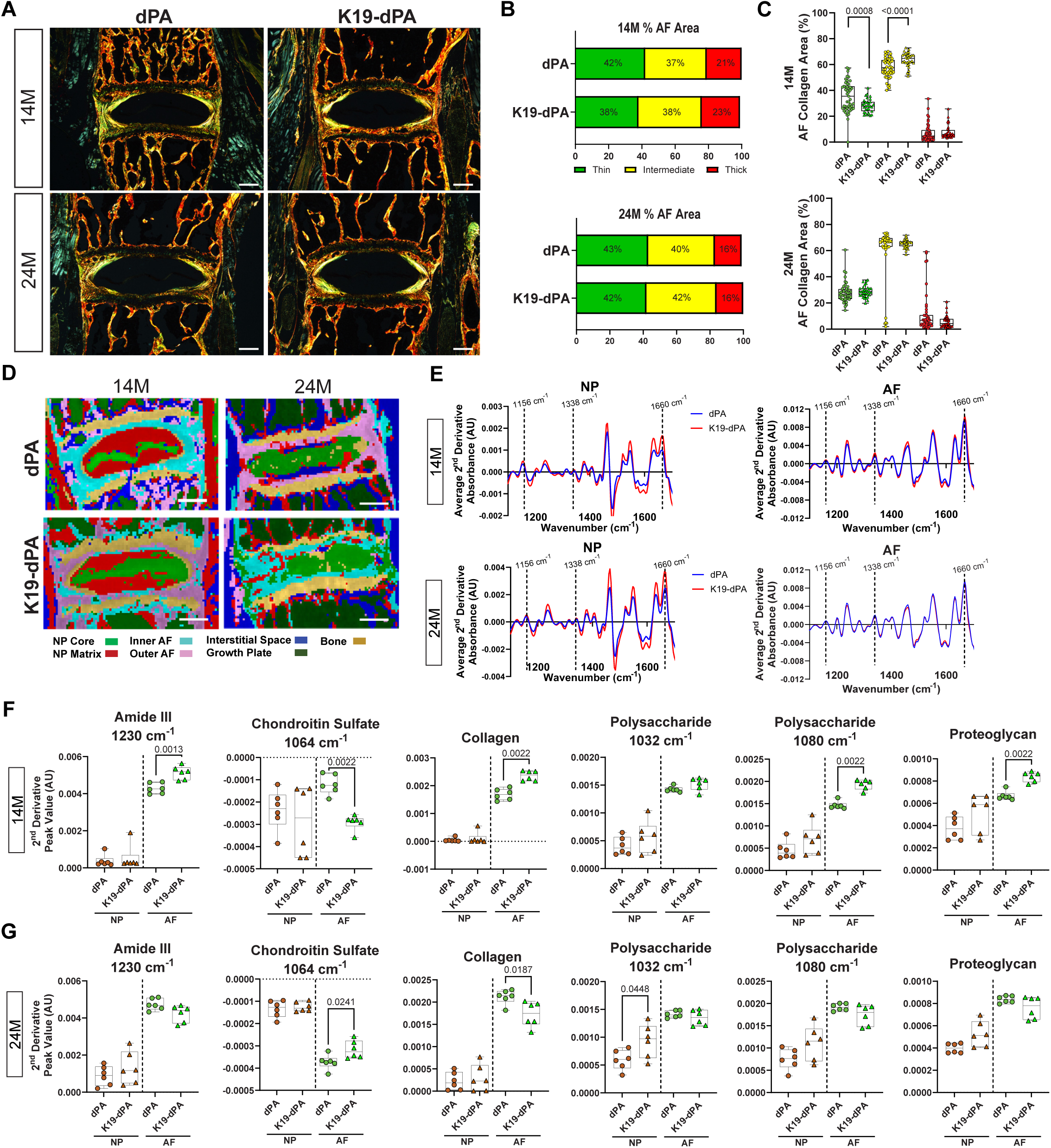
K19-dPA mice evidence altered collagen fibrillar composition and compositional changes in NP and AF. (A-C) Representative polarized images of Picrosirius Red-stained lumbar disc sections (A) and quantification of collagen fiber thicknesses (B-C) performed on 14M and 24M dPA and K19-dPA mice (scale bar in A = 200LJμm). 14M: *n* = 11 dPA, 7 K19-dPA mice, 4-5 lumbar discs/mouse, 33-48 lumbar discs/genotype and 24M: *n* = 9 dPA, 7 K19-dPA mice, 4-5 lumbar discs/mouse, 33-51 lumbar discs/genotype. (D) Spectral cluster analysis images of 14LJM and 24LJM discs (scale bar = 200LJμm). (E) Average superimposed second derivative spectra, inverted for positive visualization of the NP and AF of 14M and 24M dPA and K19-dPA mice. 14M (F) and 24M (G) quantification of mean second derivative peaks associated with chondroitin sulfate (1064LJcm^−1^), polysaccharide-associated peak (1080 cm^-1^), amide III (1230 cm^-1^), collagen (1338LJcm^−1^), and polysaccharide associated peak (1032 cm^-1^). AU = arbitrary units. For both 14M and 24M, *n* = 6 mice/genotype, 3 discs/mouse, 18 total discs/genotype. Significance for quantitative measures was determined by using Mann–Whitney *U* test or an unpaired t-test with Welch’s correction, as appropriate. Quantitative measurements represent the median with the interquartile range.

Since there were changes in AF collagen fiber content, Imaging-Fourier transform infrared (FTIR) spectroscopy was used to assess matrix compositional changes. K-means clustering was used to define broader anatomical regions of the disc using chemical compositions. The K-means clustering showed similarly defined regions between dPA and K19-dPA at 14M. However, moderate changes in the NP compartment composition were noted at 24M, suggesting the broader changes in the intervertebral disc chemical composition (Fig. 3D). Interestingly, while NP and AF spectra showed a clear and expected distinction in both genotypes, in K19-dPA mice, more prominently at 24M, AF absorbance peaks between 1400-1600 cm^-1^ showed some congruence with NP absorbance peaks within the same range, indicating diminished compositional distinction between the tissue compartments (Suppl. Fig. 1A). When the average spectra were compared between the genotypes, apparent differences in NP absorbance peaks were noticeable in K19-dPA at both 14M and 24M, suggesting there were compositional differences in the NP (Fig. 3E). Differences in the absorbance peaks of the AF between genotypes were less noticeable at both ages than NP (Fig. 3E). These results indicated that there were molecular changes in overall NP composition in K19-dPA mice.

To evaluate the specific chemical compositional differences in the NP, and AF, peaks at 1064 cm^−1^, 1120 cm^−1^, 1228 cm^−1^ (chondroitin sulfate), 1338 cm^−1^ (collagens), 1660 cm^−1^ (total protein, Amide I), 1550 cm^−1^ (Amide II), 1230 cm^−1^ (Amide III), 1032 cm^−1^ and 1080 cm^−1^ (polysaccharides), and 1156 cm^−1^ (cell-associated proteoglycans) were assessed. At 14M in K19-dPA mice, there was a higher amount of collagen and proteoglycan-associated peaks in the AF (Fig. 3D). In addition, we assessed a CS-associated peak at 1064 cm^−1^, which showed an increase in the AF (Fig. 3F). There was also an increase in the polysaccharide-associated peak at 1080 cm^−1^ and amide III-associated peak at 1230 cm^−1^ in the AF (Fig. 3F). Whereas, at 24M, there were decreases in the collagen-associated peak and an increase in the CS-associated peak 1064 cm^−1^ in the AF (Fig. 3G). There was also an increase in polysaccharide-associated peak at 1032 cm^−1^ (Fig. 3G). However, there were no differences in amide II-, chondroitin sulfate-associated peaks at 1120 cm^−1^ and 1128 cm^−1^, proteoglycan-, polysaccharide associated peak 1080 cm^−1^, and total protein-associated peaks (Suppl. Fig. 1B-C). Together the FTIR data indicate that elevated HIF-2α activity in the NP alters the molecular composition of the disc, which may result in compromised disc matrix functionality and precipitate overall disc degeneration.

### Discs of K19-dPA mice show changes in aggrecan and chondroitin sulfate abundance without affecting NP cell notochordal phenotype

To understand if the degenerative phenotype in the K19-dPA was due to changes in NP cell phenotype and major ECM constituents of the disc, we studied their expression pattern and abundance. The abundance of carbonic anhydrase 3 (CA3), an NP phenotypic marker, was comparable between the genotypes at either time point, suggesting that the NP cells did not lose their notochordal phenotype (Fig. 4A-B). Interestingly, increased ACAN levels were observed in the NP of 14M K19-dPA relative to dPA; however, this difference was not evident at 24M. Interestingly, despite increased ACAN, CS levels were lower in the NP compartment of 14M K19-dPA mice, an observation consistent with the FTIR findings and implying decreased ACAN substitution. Similarly, consistent with the increased AF proteoglycan and chondroitin sulfate (CS) content seen in FTIR analysis at the 24M, ACAN and CS levels in AF showed increased abundance in K19-dPA at 24M and levels increased as K19-dPA mice aged from 14 to 24 months suggesting transition into more mucinous matrix. On the other hand, ARGxx, which measures ADAMTS-dependent aggrecan degradation, showed no changes between genotypes at both time points. Additionally, K19-dPA mice did not show an age-dependent decrease in the abundance of cartilage oligomeric matrix protein (COMP) noted in the NP compartment of dPA control mice (Fig. 4C-D). COL I and COL II in AF also showed negligible changes with a slight increase noted in COLI abundance in the NP of 14M K19-dPA mice. These analyses suggested that while the NP notochordal phenotype is maintained, there were alterations in a few major ECM components of the disc following increased HIF-2α activity.

**Figure 4.**
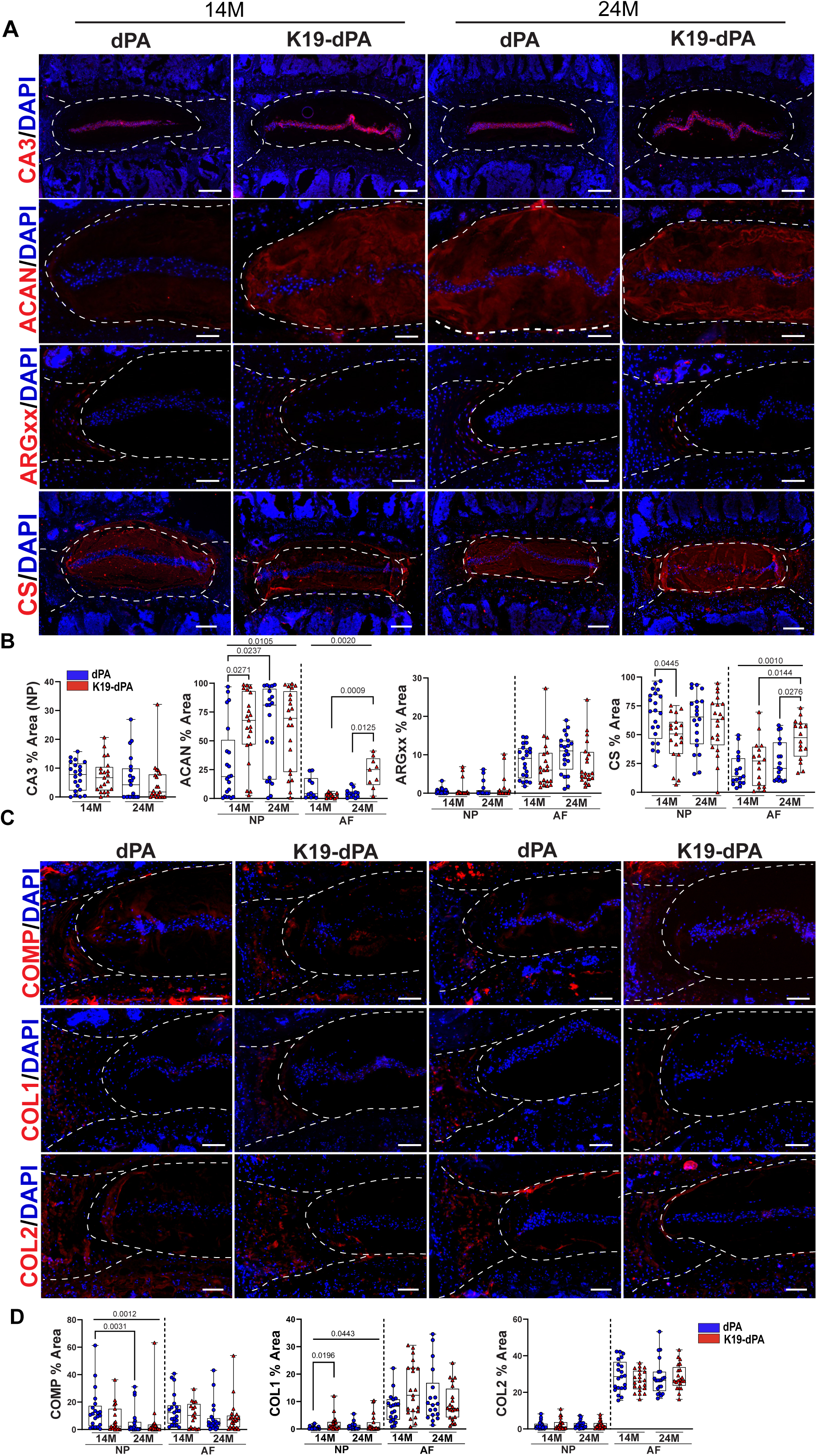
HIF-2α overexpression alters the abundance of select key ECM molecules of the disc. (A-B) Quantitative immunofluorescent staining of 14M and 24M dPA and K19-dPA lumbar discs for carbonic anhydrase 3 (CA3), aggrecan (ACAN), aggrecan neoepitope (ARGxx), and chondroitin sulfate (CS), scale bar CA3, CS = 200LJμm, scale bar ACAN, ARGxx = 100LJμm. (C-D) Quantitative immunofluorescent staining of 14M and 24M dPA and K19-dPA lumbar discs for cartilage oligomeric matrix protein (COMP), collagen I (COL1), and collagen II (COL2), scale bar = 100LJμm, *n* = 7 mice/genotype/time point, 1–3 discs/mouse, 7–21 discs/genotype/marker. White dotted lines demarcate disc compartments. Quantitative measurements represent the median with the interquartile range. Significance was determined using Kruskal–Wallis with Dunn’s test.

### Increased HIF-2**α** levels alter the transcriptomic program of the NP tissue with aging

To evaluate the global transcriptomic changes in the NP, microarray analysis was used to analyze NP tissue RNA from 14- and 24M K19-dPA and control dPA mice. Principal component analysis (PCA) showed that the 14M dPA and K19-dPA clustered separately (Fig. 5A). Hierarchical clustering of the DEGs (p ≤ 0.05, fold change (FC) ≥±1.7) was performed; this data is represented as heatmaps and volcano plots (Fig. 5B-C). To better interpret these changes, the CompBio tool was used to understand the biological context of up and downregulated DEGs, which showed super clusters comprised of top themes relating to cilia, cell motility, IGF activity regulation, hypoxia, ROBO/SLIT, and neurotrophin TRKB-receptor activity in the upregulated 14M dataset (Fig. 5D, E). The enriched DEGs in these upregulated themes included, *Dnah11, Hk2, Igfbp5, Nov and Ntrk2* (Fig. 5E, Suppl. Fig 2). One super cluster presented in the down-regulated dataset contains themes related to lipid metabolism and GLP-1 regulation with decreased expression in genes such as *Glp1*, *Adcy3*, *Fxyd3, Ppargc1, Rps6ka2* (Fig. 5D, E).

**Figure 5.**
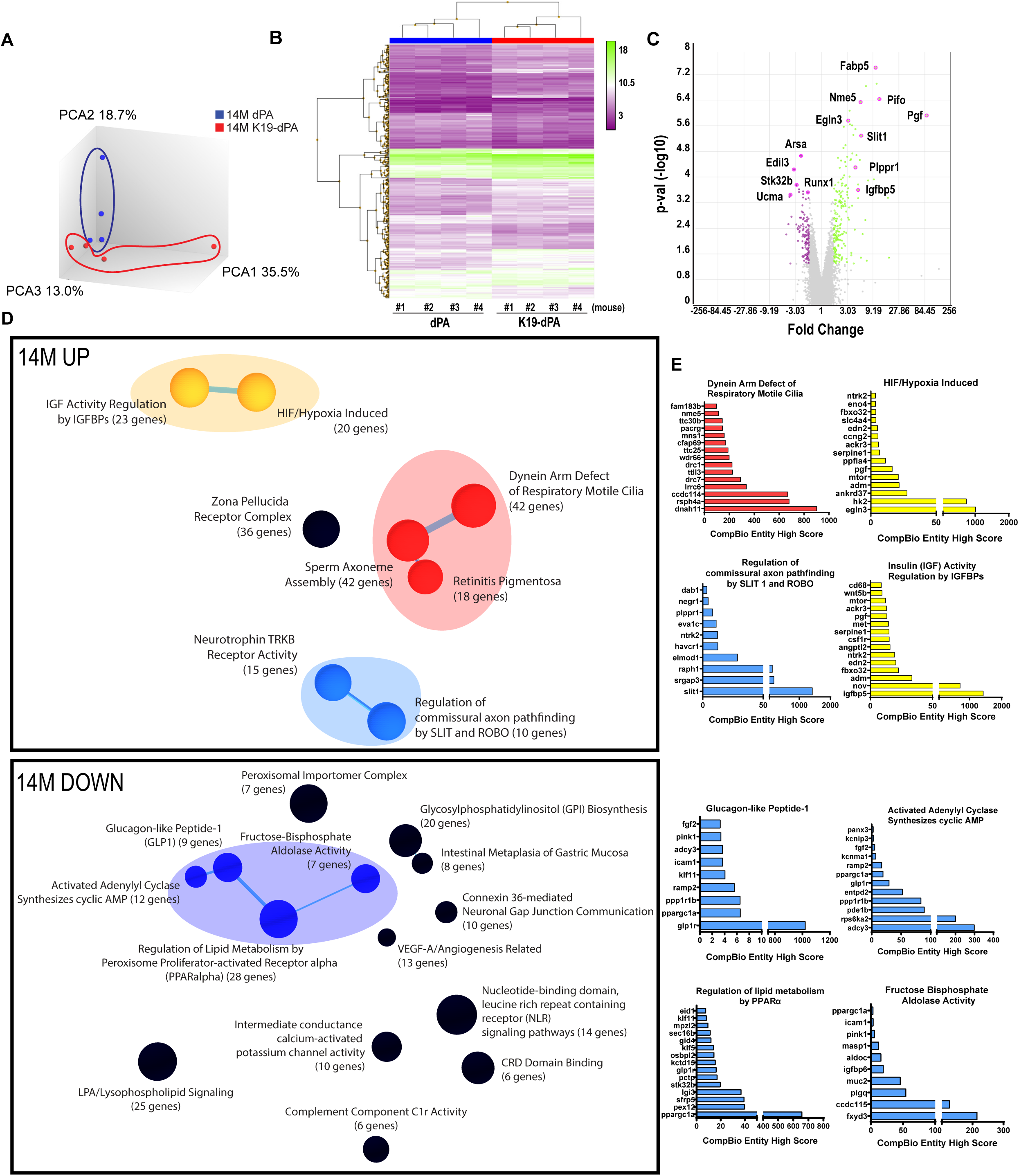
Transcriptomic analysis shows alterations in several key biological pathways in NP tissue of 14M K19-dPA mice. (A) Three-dimensional Principal component analysis (PCA) showing discrete clustering of mice based on their genotypes, *n* = 4 mice/genotype (B) Heat map and hierarchical clustering of Z-score of significantly differentially expressed genes (DEGs) from 14M K19-dPA vs dPA (p ≤ 0.05, ≥ 1.7-fold change). (C) Log-log volcano plot of DEGs in the NP shows statistical significance (P-value) versus magnitude of change (fold change). (D) CompBio analysis of DEGs and associated upregulated and downregulated themes in a ball and stick model. The enrichment of themes is shown by the size of the ball and connectedness is shown based on thickness of the lines between them. Themes of interest are colored, and superclusters comprised of related themes are highlighted. (E) Top thematic DEGs plotted based on CompBio entity enrichment score.

At 24M, PCA plots showed a clear separation between control dPA and K19-dPA samples (Fig. 6A). Identified DEGs were also plotted as heatmap and volcano plots (Fig. 6B, C) and there were more DEGs compared to seen at 14M (Fig. 6C). Within the upregulated DEGs, CompBio analysis showed unique superclusters containing themes related to ubiquitin-like hydrolase activity and SCF-mediated degradation of p27/p21 with increased expression of *Ufl1*, *Cul1*, *Skp2, Fbxw11,* and *Fbxl5* (Fig. 6D, E, Suppl. Fig. 3A). Of interest, a select group of enriched DEGs clustered into a theme named “Oxygen-Dependent Proline Hydroxylation of Hypoxia-inducible Factor Alpha” with increased expression shown in *Epas1, Cul2, and Sdha* (Fig. 6D, E). On the other hand, downregulated DEGs showed a supercluster containing themes related to actin dynamics, endodeoxyribonuclease activity, and pyrophosphatase activity with decreased expression shown in *Apex, Fen1, Sun2,* and *Ankrd2*. Interestingly, themes named “TCA Cycle” and “Weak Cry” were also found to be downregulated in 24M and contained genes including *Asns, Mdh2, Slc16a3,* and *Ror*α amongst the top genes within these themes, respectively (Fig. 6D, E, Suppl. Fig. 3B).

**Figure 6.**
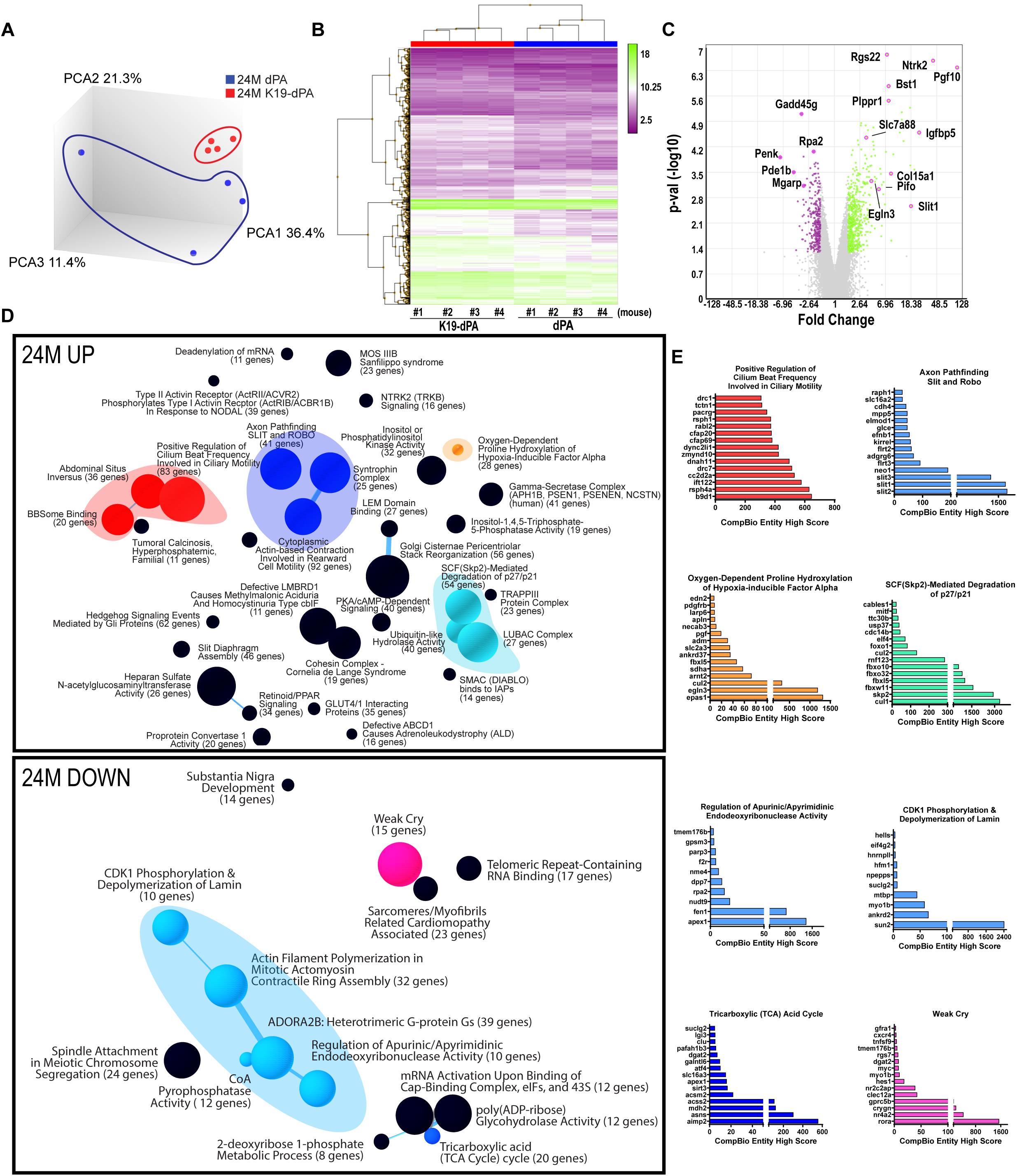
Transcriptomic analysis of NP tissues shows alterations in important biological pathways in 24M K19-dPA mice. (A) Three-dimensional Principal component analysis (PCA) showing discrete clustering of mice based on their genotypes, *n* = 4 mice/genotype (B) Heat map and hierarchical clustering of Z-score of significantly differentially expressed genes (DEGs) from 14M K19-dPA vs dPA (p ≤ 0.05, ≥ 1.7-fold change). (C) Log-log volcano plot of DEGs in the NP shows statistical significance (P-value) versus magnitude of change (fold change). (D) CompBio analysis of DEGs and associated upregulated and downregulated themes in a ball and stick model. The enrichment of themes is shown by the size of the ball and connectedness is shown based on thickness of the lines between them. Themes of interest are colored, and superclusters comprised of related themes are highlighted. (E) Top thematic up and downregulated DEGs plotted based on CompBio entity enrichment score.

Notably, at 14M and 24M, in the upregulated DEGs there was overlap with the thematic superclusters related to motile cilia, HIF/hypoxia, and SLIT/ROBO signaling. Motile cilia themes contained upregulated genes including *Dnah11, Rsph4a, Cfap69,* and *Pacrg.* The HIF/hypoxia themes had increased expression of *Egln3, Pgf, Ankrd37, Adm, and Edn2.* Whereas, the themes SLIT/ROBO signaling included common entities: *Slit1, Raph1,* and *Elmod1.* When 14M and 24M datasets were further analyzed to evaluate highly significant common genes irrespective of their thematic clustering (FDR ≤ 0.05, FC ≥ ±2.0), 11 genes including *Pgf*, *Ntrk2*, *Pifo*, *Sez6l*, *Fabp5*, *Podn*, *Slit1*, *Nme5*, *Elmod1*, *Drc1* and *Plppr1* were noted (Suppl. Fig. 4). These common DEGs are plausible HIF-2 target genes responsive to elevated HIF-2α activity, however, this hypothesis requires additional functional validation outside the scope of the present investigation. Overall, these studies strongly suggest that increased HIF-2α levels in NP alters several key biological pathways linked to cilium, cell motility, and metabolic functions.

Together our studies clearly provide evidence that elevated HIF-2α levels in the NP are pathogenic and alter many key parameters of disc health in the aging spine (Fig. 7)

**Figure 7.**
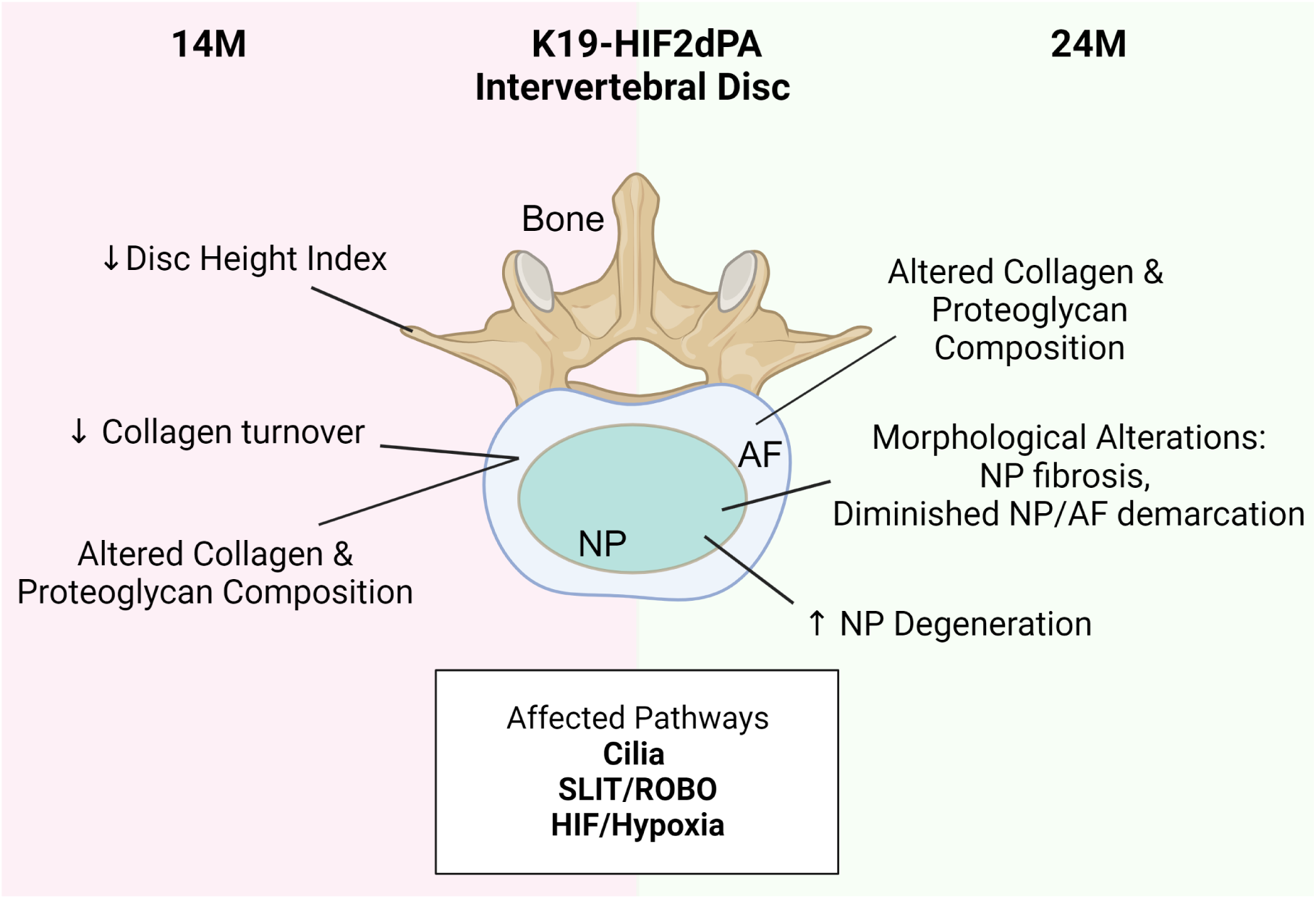
Schematic summarizing the consequences of elevated HIF-2α levels in the NP compartment on intervertebral disc health with spine aging. Created with BioRender.com.

## DISCUSSION

This study provides clear evidence that elevated HIF-2α activity in the NP causes age-dependent, mild degenerative changes in the disc corroborating our previous investigation that showed transient protection from age-related disc degeneration in mice with NP-specific deletion of HIF-2α (14). Importantly, these results underscore a possible causal link between elevated HIF-2α levels and disc degeneration seen in humans (16). It is important to note that while HIF-2α is expressed in the NP throughout the post-natal life and appears to be dispensable for maintaining the overall health of NP cells but rather pathogenic in the context of spine aging (14) It may still play other regulatory functions under specific circumstances. For example, as is noted in the growth plate and articular chondrocytes, HIF-2 may contribute to NP cell differentiation/hypertrophy or cell death in the context of disc injury, which we have not addressed in the present study (18,25,43).

The histological scoring of the NP compartment of 24-month-old K19-dPA mice showed a small but significant increase in scores compared to their littermate control mice. While the grading scores did not differ significantly, a significant decrease in disc height index (DHI) was noted at 14 months suggesting that the degenerative cascade and changes in the water-binding ability of the disc matrix were initiated earlier. This conclusion was supported by the alterations noted in the overall composition of the K19-dPA intervertebral discs. Supporting the moderate nature of phenotypic changes in the disc resulting from changes in HIF-2α levels, similar changes in the tissue morphology and matrix composition have been noted in the limb bud mesenchyme as well as the NP following the loss of HIF-2α (14,44).

The transcriptomic analysis of the NP tissue provided further insights into the pathways that were affected by the elevated HIF-2α activity in the NP and underscored the degenerative phenotype. The findings indicated that HIF-2α in the NP may upregulate the glycolytic enzyme *Hk2* at 14 months, which hints at a plausible role in HIF-2α affecting metabolic aspects, a finding that was implicated in a recent loss-of-function HIF-2α study (14). Although HIF-1α is known to play the central role in regulating glycolytic metabolism, HIF-2α has been shown to have a unique metabolic role with increased activity of this transcription factor causing downregulation of lipid metabolism and glucagon regulation at 14 months. Notably, *Glp1,* glucagon-like peptide 1, is also shown to have strong anti-inflammatory functions in many tissues (45). Importantly, GLP-1R is expressed in cartilage (46) and its agonist liraglutide is shown to exert beneficial effects on joint tissue health in the context of osteoarthritis (47). Therefore, downregulation of *Glp1* likely indicates perturbations in tissue-level anti-inflammatory mechanisms in K19-dPA mice and may in part underscore progressive degenerative changes.

The transcriptomic analysis at 24M provided further insights that metabolic alterations may underlie the observed phenotypic changes. Of interest, there was a supercluster containing one upregulated theme related to “oxygen-dependent hydroxylation of HIF-α”. This theme contained increased expression of *Epas1,* which encodes for HIF-2α, *Egln3,* and *Sdha*, succinate dehydrogenase complex A, suggesting increased mitochondrial oxidation and ROS generation in 24M NP of K19-dPA, likely contributing to the mild degenerative phenotype. Noteworthy, our lab previously showed that HIF-1α-deficient cells decreased TCA cycle flux to succinate, and lessened SDH activity, in an effort to increase flux to glutamate to maintain redox balance in NP cells (12). Whereas increased, *Egln3,* or PHD3, could implicate impaired oxygen sensing mechanisms, as this protein serves as co-activator of HIF-1 and loss-of-function of *Egln3* in mice has been shown to promote disc degeneration (48). In terms of downregulated superclusters, there was one theme contained in a cluster related to the TCA cycle with top entities including *Asna,* asparagine synthetase, which can modulate mitochondrial response (49), and decreased *Mdh2,* malate dehydrogenase 2, suggesting less formation of oxaloacetate and slowed TCA cycle activity. This theme also highlighted the downregulation of *Slc16a3,* or MCT4, a plasma-membrane monocarboxylate which is essential to maintain a balance between glycolytic and TCA cycle flux as its inhibition leads to disc degeneration (50). In this context, perhaps HIF-2α may indirectly regulate NP cell metabolism and transport mechanisms, as previously implicated for glucose and sodium-dependent transport in our HIF-2α loss-of-function *HIF-2*α*^FoxA2Cre^* mouse model (14).

Interestingly, there was another downregulated theme that contained the entity *Ror*α, RAR-related orphan receptor alpha, known to be a major circadian clock gene. We have shown that there is an interaction between BMAL1 and RORα and they modulate HIF-1α activity as well as ECM homeostasis and perturbation in circadian clock genes results in disc degeneration(51–54). Notably, corroborating our current findings, *HIF-2*α*^FoxA2Cre^* mouse has shown upregulation in circadian clock pathways. These results suggested that perturbations of cell metabolism and circadian clock may partially underscore matrix changes and degeneration in K19-HIF-2dPA mice.

Although the 14M and 24M transcriptomic data showed some unique differences, there were commonalities in their transcriptional signatures including themes related to cilia, cell motility, hypoxia, and SLIT/ROBO signaling, which were all upregulated. Our lab has shown that primary cilia alter their length in response to changes in extracellular osmolarity in the NP, thus suggesting a relationship between water-binding matrix molecules and ciliary function (55). Moreover, in response to mechanical loading, cilia are shown to maintain PTH-PTHR1 signaling axis in the NP cells to promote TGFβ1 signaling and aggrecan levels (56). Relevant to our findings in K19-dPA mice, HIF-2α in murine neuronal cells is shown to increase ciliary length in hypoxia by interacting with intraflagellar transport protein 88 homolog (IFT88) (57). Upregulation in the SLIT/ROBO cluster presented increased expression of *Slit1* which regulates secreted glycoproteins that bind to ROBO receptors known to mediate cell migration (58). An upregulation of SLIT-ROBO signaling was also observed within the NP after BNIP3 knockdown which showed dysregulation of metabolic function (59).

Identification of highly enriched common DEGs at 14- and 24-months, uncovered additional biological insights. The placental growth factor (*Pgf*) was the most upregulated common DEG, this gene is known to be expressed by ischemic or damaged tissues and has pro-angiogenic functions (60,61). It was therefore not unreasonable to assume that NP cells experienced stress from HIF-2α overexpression and *Pgf* was a compensatory response to mitigate this stress. Similarly, neurotrophic receptor tyrosine kinase 2, *Ntrk2,* was recently proposed as a prognostic gene for intervertebral disc degeneration, supporting its upregulation in this degenerative context (62). The identification of the HIF-2α responsive functions and genes in AF and the endplate would provide complementary information to understand the compartment-specific functionality of HIF-2α in the disc (63–65). While our studies draw clear distinctions between the non-redundant functions of two HIF-α isoforms in the disc, inhibition of HIF-2α activity in the context of spine aging is likely to provide beneficial effects in maintaining disc health.

## Data availability statement

The datasets generated and analyzed during this study are available in the NCBI Gene Expression Omnibus (GEO) repository with accession number GSE249908.

## Ethics

All mouse care procedures, housing, breeding, the collection of animal tissues, and experiments were performed under protocols approved by the Institutional Animal Care and Use Committee (IACUC) of Thomas Jefferson University following the IACUC’s relevant guidelines and regulations.

## Acknowledgments

We would like to thank Dr. Ernestina Schipani, University of Pennsylvania for providing the HIF-2αdPA allele. Research reported in this publication was supported by R01AR055655, R01AR074813, and R01 AG073349 from the National Institute of Arthritis and Musculoskeletal and Skin Diseases (NIAMS) and National Institute on Aging (NIA) of the National Institutes of Health. Shira Johnston was supported by T32 AR052273.

## Disclosures

R.A. Barve may receive royalty income based on the CompBio technology developed by R.A. Barve and licensed by Washington University to PercayAI. The remaining authors declare they have no competing interests to disclose about the contents of this article.

## Supplementary Figure Legends

**Supplementary Figure 1.** (A) NP and AF superimposed average second derivative spectra, inverted for positive visualization from 14M and 24M dPA and K19-dPA NP mice. (B) Quantification of mean second derivative peaks associated with total protein (1660 cm^−1^), cell-associated proteoglycan (1156 cm^−1^), and AU = arbitrary units. *n* = 6 mice/genotype, 3 discs/mouse, 18 total discs/genotype. Significance for quantitative measures was determined by using Mann–Whitney *U* test or an unpaired t-test with Welch’s correction, as appropriate. Quantitative measurements represent the median with the interquartile range.

**Supplementary Figure 2.** 14-month upregulated DEGs from select central themes plotted based on their CompBio entity scores. *n* = 4 mice/genotype/time point.

**Supplementary Figure 3. (A)** 24-month upregulated and (B) down-regulated DEGs from select central themes plotted based on their CompBio entity scores. *n* = 4 mice/genotype/time point.

**Supplementary Figure 4. (A)** The highly enriched common DEGs in the NP between 14M and 24M K19-dPA mice with a fold change≥ ±2 and FDR ≤ 0.05. *n* = 4 mice/genotype/time point

