## Supplementary figures and images for "Increased HIF-2α Activity in the Nucleus Pulposus Causes Intervertebral Disc Degeneration in the Aging Mouse Spine"

### Supplementary Figure 1

Supplementary Fig. 1

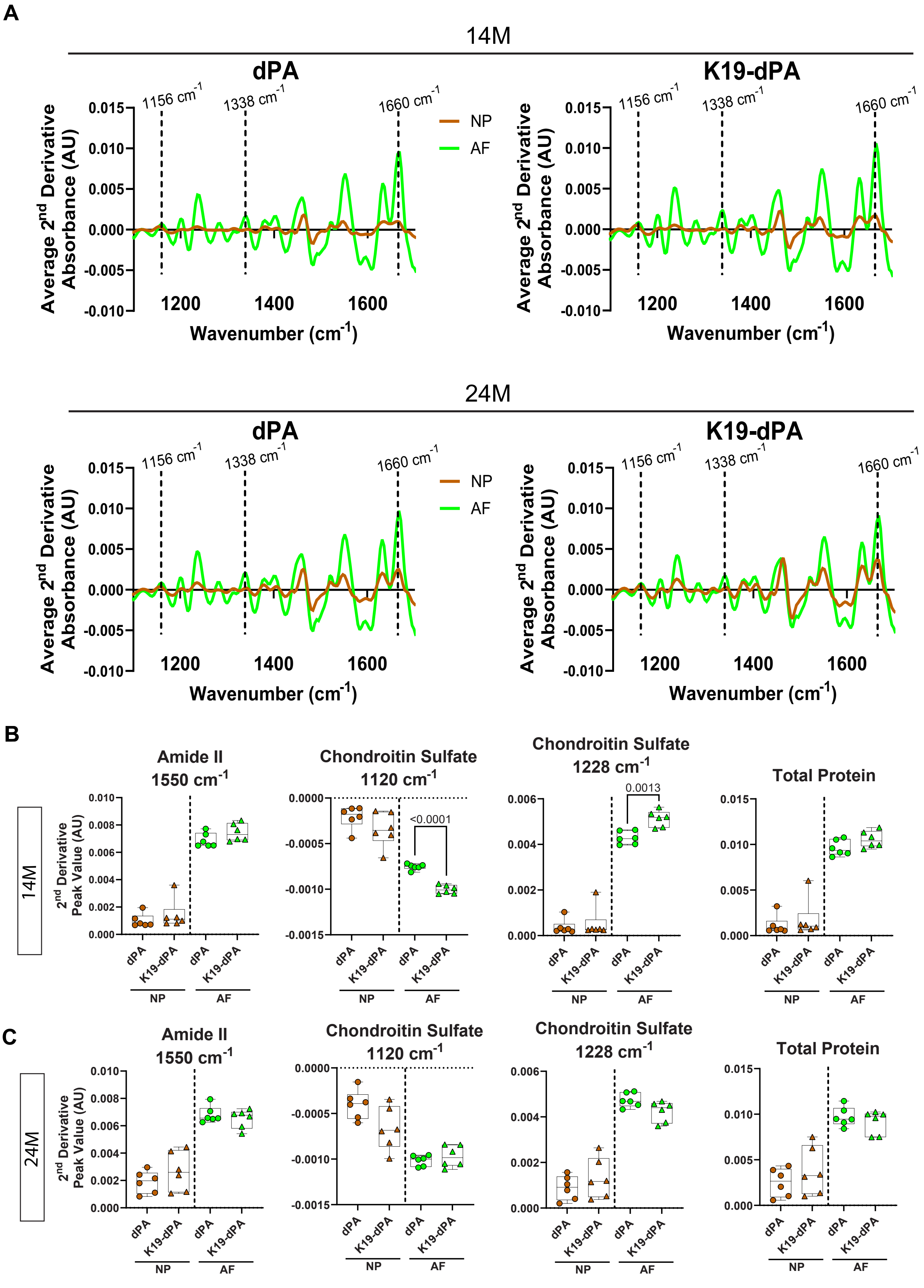

### Supplementary Figure 2

Supplementary Figure 2

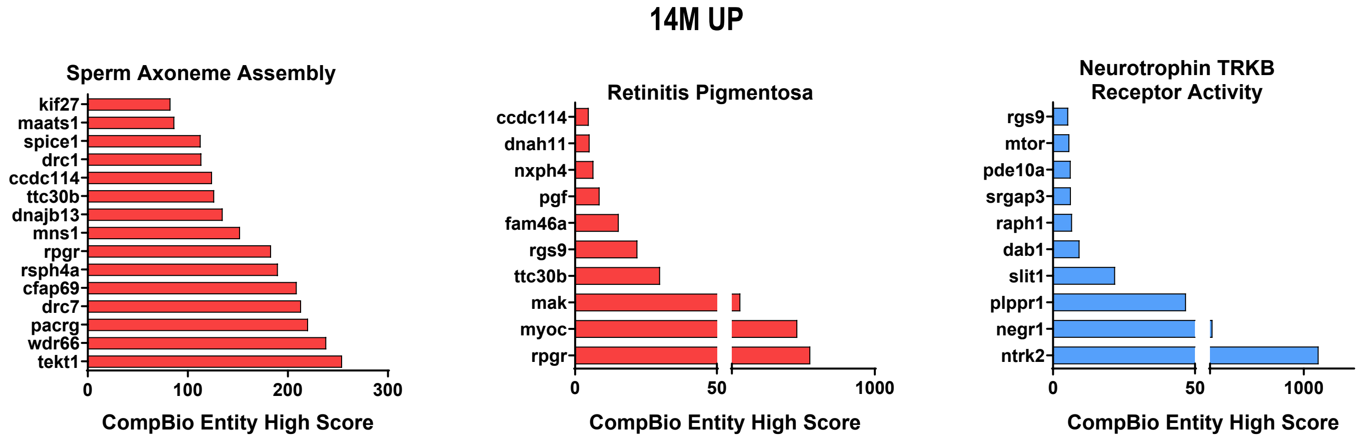

### Supplementary Figure 3

Supplementary Figure 3

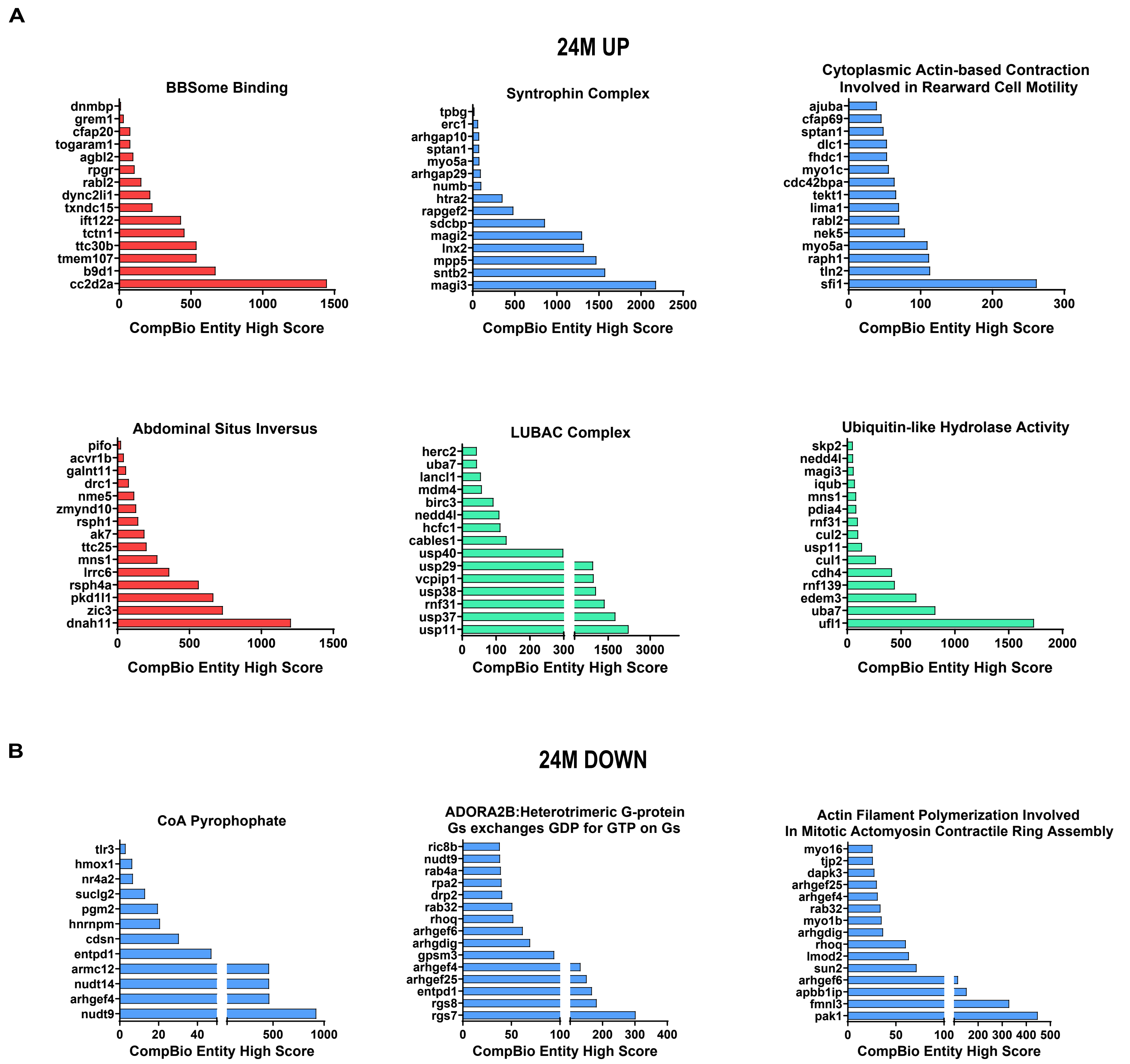

### Supplementary Figure 4

Supplementary Fig. 4

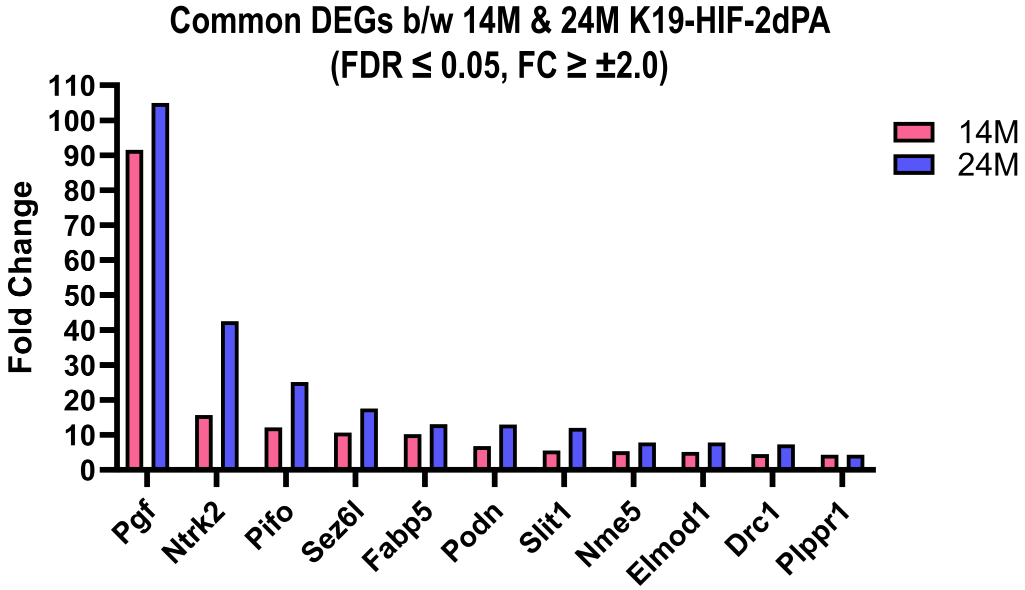
